# Naringenin Attenuates Cisplatin-Induced Hepatotoxicity and Nephrotoxicity by Restoring Glutathione Homeostasis and Suppressing Lipid Peroxidation in a Murine Model

**DOI:** 10.64898/2026.04.22.720080

**Authors:** Ashwin Dev, Kamalesh Dattaram Mumbrekar

## Abstract

Cisplatin is a cornerstone chemotherapeutic agent for a broad spectrum of solid malignancies, yet its clinical utility is substantially curtailed by dose-limiting organ toxicity, principally nephrotoxicity and hepatotoxicity, mediated through reactive oxygen species (ROS)-driven oxidative stress, glutathione depletion, and lipid peroxidation. Naringenin (NAR), a bioactive citrus flavanone, possesses potent free-radical scavenging, anti-inflammatory, and cytoprotective properties that make it a compelling candidate for chemoprotection. The present study investigated whether oral naringenin supplementation (50 mg/kg body weight/day for 30 days) could mitigate cisplatin-induced oxidative injury to the liver and kidney in male Swiss albino mice. Cisplatin was administered intraperitoneally at 2.3 mg/kg body weight in three cycles of five consecutive days followed by a five-day interval. Biochemical indices of oxidative stress, such as malondialdehyde (MDA), reduced glutathione (GSH), and glutathione S-transferase (GST) activity, were assayed in liver and kidney homogenates on day 45. Cisplatin administration significantly elevated hepatic and renal MDA levels, indicating pronounced lipid peroxidation, and markedly depleted the concentrations of GSH and the activity of GST in both organs. Compared with cisplatin alone, naringenin coadministration significantly attenuated the increase in the level of MDA, restored the level of GSH, and rescued the activity of GST in both tissues, with more pronounced effects in the kidney. Notably, compared with the control, naringenin alone did not alter any biochemical parameters, confirming its physiological safety at the administered dose. These findings demonstrate that naringenin has meaningful hepatoprotective and nephroprotective effects against cisplatin-induced oxidative toxicity, possibly through antioxidant augmentation, glutathione repletion, and membrane stabilization mechanisms. This study provides a rational preclinical basis for evaluating naringenin as a coadministered chemoprotectant in cisplatin-based chemotherapy regimens.

## Introduction

Cancer remains among the foremost causes of morbidity and mortality worldwide, accounting for an estimated 19.3 million new cases and 10.0 million deaths globally in 2020 [1]. Chemotherapy constitutes a central pillar of systemic cancer treatment, either as a definitive modality or in combination with surgery and radiation. Among the arsenal of cytotoxic agents, cisplatin (cis-diaminedichloroplatinum II; CDDP) occupies a preeminent position and is widely regarded as one of the most effective broad-spectrum antineoplastic drugs available. Since its serendipitous discovery by Rosenberg et al. in 1965 and subsequent clinical introduction in the 1970s, cisplatin has remained a first-line treatment for testicular, ovarian, cervical, head and neck, bladder, esophageal, lung, and brain malignancies, as well as mesothelioma [2, 3, 4].

The antineoplastic mechanism of cisplatin is principally attributable to its ability to form intra- and interstrand crosslinks with genomic DNA after it is acquired in the intracellular milieu. Following cellular uptake via the copper transporter CTR1 and, to a lesser extent, organic cation transporters, cisplatin undergoes sequential chloride dissociation in the low-chloride cytoplasmic environment, generating reactive aquated monoaquo and diaquo species that preferentially target the N7 positions of purine bases [4, 5]The resulting bifunctional Pt-DNA adducts distort the double helix, impede DNA replication, and trigger cell cycle arrest and apoptosis through the activation of p53, downstream caspase cascades, and the intrinsic mitochondrial apoptotic pathway [6]. Concurrently, cisplatin activates p53 and alters the transcription of PPAR-α and its coactivator PGC-1α, with a downstream reduction in medium-chain acyl dehydrogenase (MCAD) and upregulation of pyruvate dehydrogenase kinase isozyme 4 (PDK4), reflecting profound disruption of mitochondrial fatty acid oxidation and energy metabolism [7].

Despite its unequivocal clinical efficacy, the therapeutic utility of cisplatin is significantly constrained by a constellation of dose-limiting toxicities that affect multiple organ systems. Common acute adverse effects include nausea, emesis, myelosuppression, alopecia, and peripheral neuropathy [8, 9]. However, the most clinically consequential organ-specific toxicities are nephrotoxicity and hepatotoxicity, which are frequently cumulative, dose dependent, and potentially irreversible.

Nephrotoxicity is the most widely documented and dose-limiting toxicity of cisplatin and affects approximately 25–35% of patients receiving standard-dose regimens [10]. Compared with plasma, the kidney accumulates five-to tenfold more cisplatin, predominantly in the proximal tubule, via active uptake through organic cation transporter 2 (OCT2) [11]. At the renal tubular level, cisplatin depletes antioxidant defenses, principally glutathione and catalase, in the renal cortex, generating superoxide anions, hydrogen peroxide, and highly reactive hydroxyl radicals [12, 13]. The resulting oxidative milieu drives tubular apoptosis and necrosis, which manifests clinically as a reduction in the glomerular filtration rate (GFR), tubular dysfunction, electrolyte imbalance (hypomagnesemia, hypokalemia), and elevated serum creatinine and blood urea nitrogen (BUN), which are hallmarks of acute kidney injury (AKI) [14]. Mechanistically, cisplatin also inhibits Na+/K+-ATPase by binding to its cytoplasmic domains, disrupting tubular electrolyte reabsorption and exacerbating mitochondrial dysfunction via impairment of electron transport chain complexes [7].

Hepatotoxicity, although less prominently documented than nephrotoxicity, is an equally significant clinical concern with cisplatin therapy. The liver, as the primary organ of xenobiotic metabolism and detoxification, is exposed to high concentrations of cisplatin and its metabolites during systemic distribution. Cisplatin-induced hepatic injury is characterized by elevated serum transaminases (ALT and AST), alkaline phosphatase, and bilirubin, reflecting hepatocellular necrosis, cholestasis, and impaired hepatic synthetic function [15]. At the molecular level, cisplatin depletes hepatic GSH reserves, inhibits the activity of glutathione-S-transferase (GST), a key phase II detoxification enzyme, and promotes lipid peroxidation via ROS-mediated oxidation of polyunsaturated fatty acids (PUFAs) in hepatocyte membranes, yielding malondialdehyde (MDA) as the principal biomarker [12, 13]. Furthermore, cisplatin upregulates heme oxygenase-1 (HO-1) expression in the liver as part of a compensatory, although ultimately insufficient, antioxidant response. The hepatic glutathione system, which comprises GSH, GST, glutathione reductase, and glutathione peroxidase, is the central enzyme involved in the defense against cisplatin-induced oxidative hepatocellular damage, and its depletion markedly amplifies cellular vulnerability to ROS-mediated injury.

The urgent need to develop effective chemoprotective strategies that preserve organ function without compromising the antitumor efficacy of cisplatin has prompted extensive investigations of naturally derived phytochemicals. Naringenin (4′,5,7-trihydroxyflavanone; NAR) is a bioactive flavanone that is abundant in citrus fruits, including grapefruits, oranges, lemons, and tangerines, as well as in tomatoes and certain herbs [16]. Structurally, naringenin possesses a flavanone backbone with three hydroxyl groups at positions 4, 5, and 7, which confer its potent electron-donating antioxidant capacity and ability to form stable complexes with metal ions, including platinum [17].

Pharmacologically, naringenin exhibits a diverse repertoire of biological activities, including free-radical scavenging, anti-inflammatory, antiproliferative, anti-atherogenic, hypolipidemic, and immunomodulatory effects [18, 19, 20]. Naringenin exerts antioxidant activity through multiple complementary mechanisms: (i) direct ROS neutralization by the donation of phenolic hydroxyl groups; (ii) intercalation between phospholipid bilayers, reducing membrane fluidity and sterically protecting PUFAs from oxidative attack; (iii) modulation of cytochrome P450-dependent monooxygenases (CYP1A1 and CYP1B1) involved in xenobiotic activation and detoxification; and (iv) upregulation of endogenous antioxidant enzymes via Nrf2/ARE pathway activation [17, 21, 22]. Naringenin has demonstrated hepatoprotective efficacy in models of carbon tetrachloride-induced liver fibrosis, nonalcoholic fatty liver disease, and drug-induced hepatotoxicity and has been shown to have nephroprotective effects in diabetic and ischemia□reperfusion injury models [17, 21]. Critically, accumulating evidence indicates that naringenin does not reduce tumor platinum concentrations, suggesting that it may exert chemoprotection without compromising the antineoplastic bioavailability of cisplatin [23].

Despite this compelling mechanistic rationale, systematic in vivo evidence for the capacity of naringenin to mitigate cisplatin-induced oxidative damage in both the liver and kidney simultaneously under a clinically relevant cyclical dosing regimen has remained largely absent from the literature. The present study was therefore designed to investigate the hepatoprotective and nephroprotective potential of oral naringenin coadministration against cisplatin-induced oxidative toxicity in male Swiss albino mice, as assessed by validated biochemical markers of oxidative stress, antioxidant capacity, and glutathione metabolism. We hypothesized that naringenin’s multimodal antioxidant mechanisms would significantly attenuate cisplatin-induced MDA elevation and restore GSH and GST levels in both the liver and kidney, providing a preclinical rationale for its evaluation as a clinical chemoprotectant.

## Materials and Methods

### Animals and Ethical Statement

Male Swiss albino mice (6–8 weeks of age, body weight 30–35 g) were procured from the Central Animal Research Facility, Manipal Academy of Higher Education (MAHE), Manipal, India. The animals were housed in individually ventilated polypropylene cages (five per cage) under standardized conditions: an ambient temperature of 23 ± 2°C, a relative humidity of 50 ± 5%, and a 12-hour light/12-hour dark photoperiod. Standard rodent pelleted diet and sterilized potable water were available ad libitum throughout the study period. All experimental protocols were conducted in strict compliance with the guidelines of the Committee for the Purpose of Control and Supervision of Experiments on Animals (CPCSEA), Government of India. Ethical approval was obtained from the Institutional Animal Ethics Committee (IAEC), Kasturba Medical College, Manipal Academy of Higher Education (IAEC/KMC/15/2019). Male animals were exclusively used to avoid hormonal variability associated with the estrous cycle in female mice, which may confound biochemical endpoints. The study protocol adhered to the ARRIVE 2.0 guidelines for reporting animal experiments [24].

### Drugs and chemicals

Cisplatin (cis-diaminedichloroplatinum II; CDDP; purity ≥ 99.9%) was purchased from Sigma□Aldrich Co. LLC (St. Louis, MO, USA) and dissolved in sterile normal saline (0.9% NaCl) immediately before each administration. Naringenin (4′,5,7-trihydroxyflavanone; NAR; purity ≥ 95%) was obtained from SRL Laboratories (Mumbai, India) and suspended in sterile Milli-Q water with gentle vortexing. Reduced glutathione (GSH), 5,5′-dithiobis-(2-nitrobenzoic acid) (DTNB), 1-chloro-2,4-dinitrobenzene (CDNB), thiobarbituric acid (TBA), trichloroacetic acid (TCA), Bradford reagent, and all other analytical-grade reagents were obtained from Sigma□Aldrich or Himedia Laboratories (Mumbai, India).

### Experimental Design and Treatment Groups

Twenty-four male Swiss albino mice were randomly allocated into four groups, i.e., control, cisplatin alone, naringenin alone and a combination of cisplatin and naringenin (each group n = 6). The dose of 2.3 mg/kg body weight cisplatin was derived from allometric scaling of the minimum effective human clinical dose and is consistent with previously established murine nephrotoxicity models [25, 26]. Cisplatin was administered intraperitoneally in three identical cycles, each comprising five consecutive daily doses followed by a five-day drug-free interval, for a total of 30 days of experimental duration. Naringenin was administered orally by gavage at 50 mg/kg body weight once daily throughout the entire 30-day period, starting five days before the first cisplatin dose in the combination group to allow preloading of tissue antioxidant defenses, which is consistent with chemoprotective dosing strategies employed in the literature [21]. The animals were monitored daily for clinical signs of toxicity, behavioral changes, and body weight. All the animals survived the study endpoint. Tissue collection was performed on day 31 (15 days after the final cisplatin dose), under anesthesia followed by cervical dislocation.

### Tissue collection and preparation of homogenates

Following euthanasia, the liver and kidneys were rapidly excised, rinsed in ice-cold phosphate-buffered saline (PBS, pH 7.4), blotted dry, and weighed. Tissue aliquots were snap-frozen in liquid nitrogen and stored at −80°C until analysis. For the biochemical assays, the tissues were thawed on ice and homogenized in ice-cold PBS (pH 7.4) at a 1:20 (w/v) dilution using a high-speed homogenizer (Remi Homogenizer 8000, India). The homogenates were centrifuged at 13,500 rpm for 25 minutes at 4°C (Eppendorf 5810R), and the supernatant was collected and used for all subsequent biochemical determinations. The total protein concentration in each supernatant was quantified by the Bradford assay [27], using bovine serum albumin (BSA) as the standard, and the absorbance was measured at 595 nm on a Shimadzu spectrophotometer (Shimadzu Corporation, Japan). All biochemical assay results were normalized to milligrams of protein to permit intergroup comparisons.

### Biochemical Assays

Lipid peroxidation: Thiobarbituric acid reactive substances (TBARS) assay

Lipid peroxidation was quantified by measuring malondialdehyde (MDA) as a thiobarbituric acid reactive substance (TBARS) according to the methods of Utley et al. (1967), with minor modifications. Briefly, 500 µL of tissue homogenate was mixed with an equal volume of 10% trichloroacetic acid (TCA) and centrifuged at 3,000 rpm for 10 minutes at 4°C to precipitate proteins. Aliquots of the clarified supernatant were then mixed with 250 µL of 0.67% thiobarbituric acid (TBA) in 0.1 M HCl and incubated in a boiling water bath (90–95°C) for 30 minutes. After the samples were cooled to room temperature, the absorbance was measured at 532 nm and 600 nm using a spectrophotometer. The MDA concentration was calculated using the equation [MDA] (mM) = (A_532_ − A_600_)/155 mM^−1^ cm^−1^, where 155 mM^−1^ cm^−1^ is the molar extinction coefficient of the MDA–TBA adduct. The results are expressed as nmol MDA/mg protein [28].

### Estimation of Reduced Glutathione (GSH)

Hepatic and renal GSH concentrations were determined according to the colorimetric method of Rahman et al. (2007) on the basis of the reaction with 5,5′-dithiobis-(2-nitrobenzoic acid) (DTNB; Ellman’s reagent). Briefly, 500 µL of tissue supernatant was deproteinized by adding an equal volume of 5% sulfosalicylic acid and incubating at 4°C for 30 minutes. After centrifugation at 12,000 rpm for 5 minutes at 4°C, 250 µL of the supernatant was combined with 100 µL of 0.06 mM DTNB in phosphate buffer and 650 µL of 0.1 M sodium phosphate buffer (pH 7.4). The reaction mixture was incubated at 37°C for 30 minutes, and the absorbance was read at 412 nm. The GSH concentration was interpolated from a standard curve prepared using known concentrations of reduced glutathione (5–100 µM). The results are expressed as µmol GSH/mg protein [29].

### Glutathione S-Transferase (GST) Activity Assay

GST activity was measured spectrophotometrically using 1-chloro-2,4-dinitrobenzene (CDNB) as the substrate, according to the methods of Boyland and Chasseaud (1969). The assay cocktail contained 1 mM CDNB, 1 mM reduced glutathione, and 0.1 M sodium phosphate buffer (pH 6.5). The reaction was initiated by the addition of 50 µL of tissue supernatant to the cocktail, and the increase in absorbance at 340 nm was recorded continuously for 5 minutes at 25°C on a UV-1800 Shimadzu spectrophotometer. GST activity was calculated using the molar extinction coefficient of the GSH–CDNB conjugate (9.6 ^mM−1 cm−1^) and expressed as units (µmol substrate conjugated/minute) per milligram of protein (U/mg protein) [30].

### Statistical Analysis

All the data are presented as the mean ± standard error of the mean (SEM). Statistical comparisons between the four groups were performed using one-way analysis of variance (ANOVA) followed by a post hoc test to correct for multiple pairwise comparisons while controlling for the familywise error rate. All analyses were conducted using GraphPad Prism version 8.0 (GraphPad Software Inc., San Diego, CA, USA). A value of p < 0.05 was considered to indicate statistical significance.

## Results

### Effects on Lipid Peroxidation (MDA Levels)

The MDA concentrations in liver and kidney homogenates across the experimental groups are shown in Figure 1. Analysis of the liver (Figure 1A) revealed that compared with vehicle treatment, cisplatin treatment resulted in a significant increase in hepatic MDA levels (p = 0.0034), indicating that oxidative stress-driven lipid peroxidation occurred in liver tissue. The naringenin-alone group and the cisplatin + naringenin combination group did not differ significantly from the controls (p > 0.05 for both), suggesting that naringenin cotreatment effectively attenuated cisplatin-induced hepatic lipid peroxidation and that naringenin alone did not affect the activity at the administered dose.

**Fig 1.**
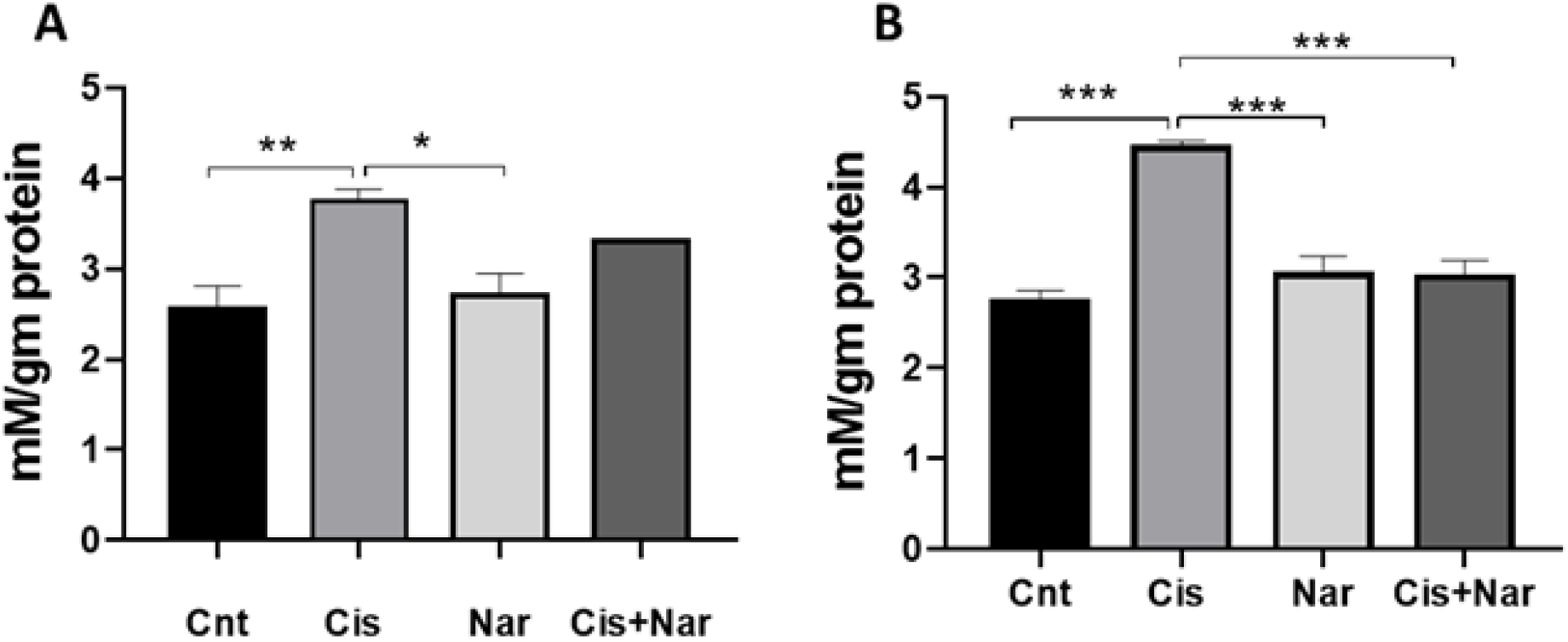
Comparison of malondialdehyde (MDA) levels (nmol/mg protein) across experimental groups. (A) Liver; (B) Kidney. The data are presented as the mean ± SEM (n = 6). Statistical significance was determined by one-way ANOVA with Tukey’s post hoc test. *p < 0.05; **p < 0.01; ***p < 0.001 relative to the indicated comparisons.

In the kidney (Figure 1B), the intergroup differences were more pronounced, with higher renal MDA levels in the cisplatin group than in the control group (p = 0.0001), which is consistent with the well-established preferential accumulation of cisplatin in the renal proximal tubule and the consequent intensification of local oxidative stress. Critically, the MDA levels were significantly lower in the cisplatin + naringenin group than in the cisplatin-alone group (p = 0.0004), and compared with the cisplatin group, the naringenin-alone group also had significantly lower MDA levels (p = 0.0058), confirming the antioxidant efficacy of naringenin in the kidney. Neither the naringenin nor the cisplatin + naringenin group differed significantly from the controls in renal MDA, underscoring the near-complete restoration of lipid peroxide homeostasis by naringenin cotreatment. Compared with the liver, the kidney exhibited markedly greater absolute increase in MDA in response to cisplatin, which is consistent with the organ’s selective vulnerability to cisplatin toxicity.

### Effect on Reduced Glutathione (GSH) Levels

In the liver (Figure 2A), comparison revealed that compared with the naringenin-alone group, the cisplatin-alone group had significantly lower GSH levels (p = 0.0340), reflecting cisplatin-mediated hepatic glutathione depletion. While the GSH level tended to be lower in the cisplatin group than in the control group, this difference did not reach statistical significance, possibly reflecting a slight effect on hepatic GSH depletion or differential susceptibility of hepatic versus renal glutathione pools to cisplatin at the administered dose. The GSH levels of the cisplatin + naringenin group were comparable to those of the control group, suggesting partial hepatic GSH restoration by naringenin. Naringenin alone did not significantly alter hepatic GSH levels compared with those in controls.

**Fig 2.**
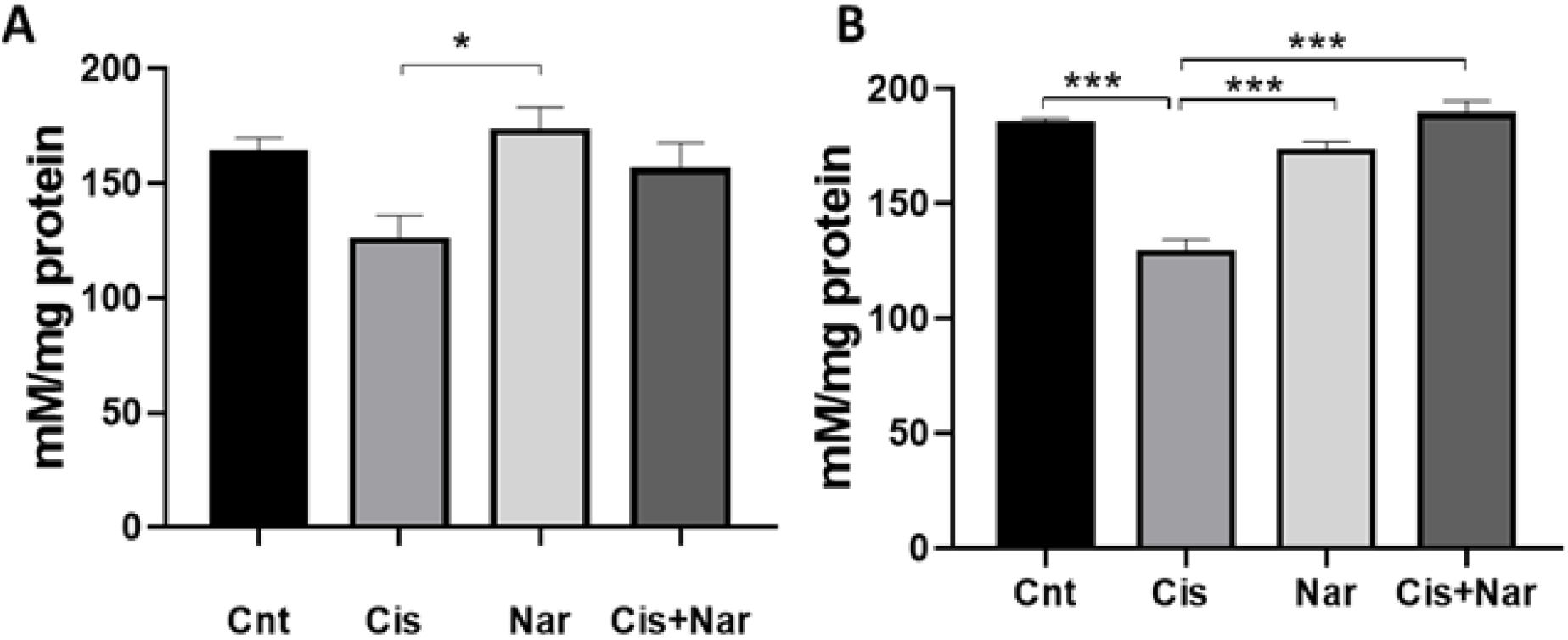
Comparison of reduced glutathione (GSH) levels (µmol/mg protein) across experimental groups. (A) Liver; (B) Kidney. Data are presented as the mean ± SEM (n = 6). *p < 0.05; **p < 0.01; ***p < 0.001 (one-way ANOVA, Tukey’s post hoc test).

In kidney tissue, compared with the control treatment, cisplatin treatment caused a profound depletion of renal GSH (p < 0.0001), corroborating the established mechanism of cisplatin-mediated inhibition of glutathione reductase activity and direct platinum-GSH conjugation, leading to net glutathione exhaustion. Renal GSH levels were significantly greater in the naringenin-alone group than in the cisplatin group (p = 0.0002), and in the cisplatin + naringenin group, this protective increase was even more pronounced (p < 0.0001 vs. cisplatin). Neither the naringenin alone nor the combination group differed significantly from the controls, indicating that naringenin cotreatment substantially preserved renal GSH reserves against cisplatin-induced depletion. The degree of renal GSH depletion substantially exceeded that in the liver, further emphasizing the kidney’s particular susceptibility to cisplatin-induced antioxidant exhaustion.

### Effect on Glutathione S-Transferase (GST) Activity

In the liver (Figure 3A), compared with the control, cisplatin markedly suppressed hepatic GST activity (p = 0.0003), reflecting the dual impact of cisplatin on phase II detoxification: depletion of the GSH substrate and direct inhibition of GST enzymatic activity by platinum-mediated modification of critical enzyme thiol residues. Compared with the cisplatin group, the naringenin-alone group maintained significantly greater GST activity (p = 0.0004), and compared with the cisplatin group, the cisplatin + naringenin combination group similarly showed significantly preserved GST activity (p = 0.0008). No significant differences were observed between the control, naringenin, or combination groups, confirming that naringenin preserved constitutive GST activity and compensated for cisplatin-mediated enzyme suppression.

**Fig 3.**
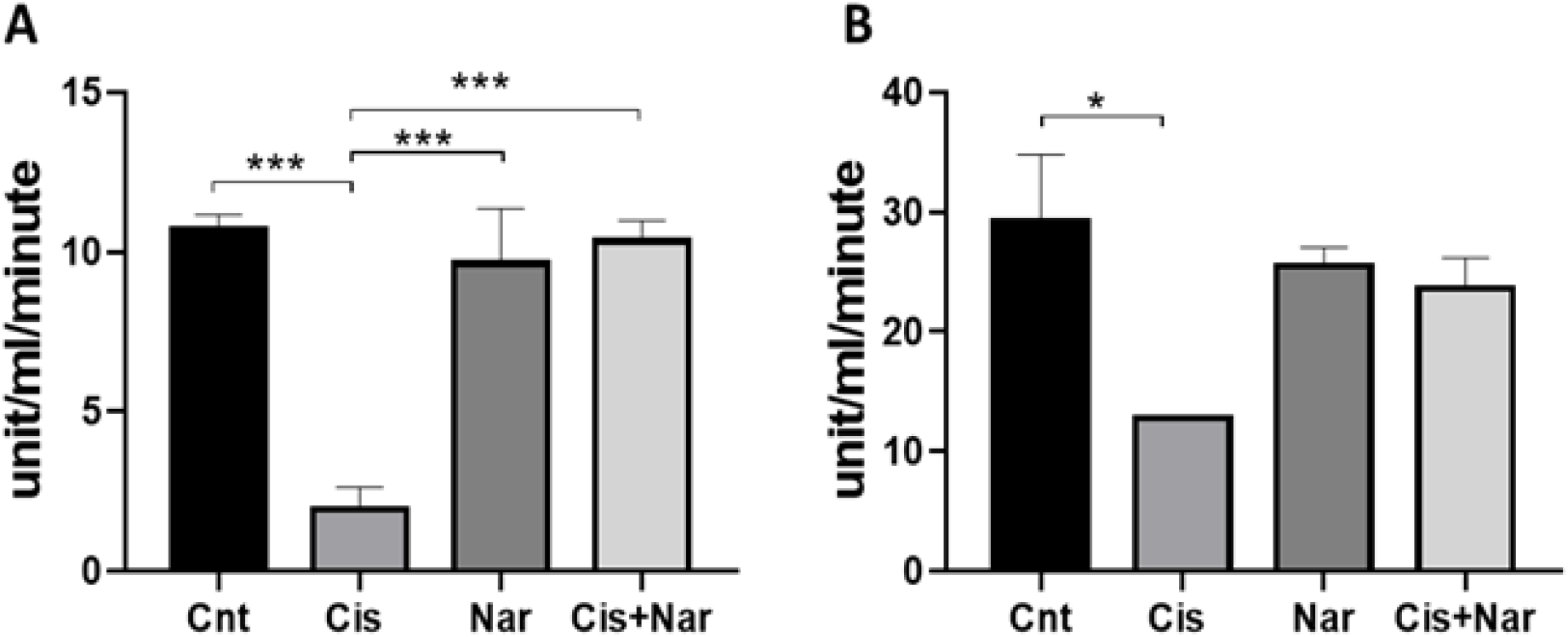
Comparison of glutathione S-transferase (GST) activity (U/mg protein) across experimental groups. (A) Liver; (B) Kidney. Data are presented as the mean ± SEM (n = 6). *p < 0.05; **p < 0.01; ***p < 0.001 (one-way ANOVA, Tukey’s post hoc test).

In the kidney (Figure 3B), renal GST activity was significantly lower in the cisplatin-treated group than in the control group (p = 0.0145). While the renal GST activity tended to increase in the naringenin and combination groups compared with the cisplatin group, these differences did not reach statistical significance (p > 0.05), suggesting that the preservation of renal GST activity by naringenin may be less pronounced than its effects on renal GSH levels and MDA or that the sample size limits the detection of subtler renal GST changes. No group differed significantly from the controls for renal GST, except for the cisplatin-alone group. Collectively, these data indicate that naringenin most potently rescues GST activity in the liver, whereas its nephroprotective effects are primarily mediated through the augmentation of GSH activity and the suppression of MDA activity.

## Discussion

The present study provides systematic preclinical evidence that oral naringenin supplementation significantly attenuates cisplatin-induced oxidative hepatotoxicity and nephrotoxicity in mice, as evidenced by a significant reduction in MDA levels, restoration of GSH levels, and preservation of GST activity in both the liver and kidney. These findings are both mechanistically coherent and clinically relevant, adding to the growing body of evidence supporting the chemoprotective potential of citrus flavanones against platinum-based chemotherapy-induced organ toxicity.

An oxidative stress model of cisplatin-induced organ toxicity is well established. Once inside the cell, the aquate derivatives of cisplatin react with intracellular GSH both spontaneously and enzymatically via GST, forming stable platinum□glutathione complexes that are subsequently exported via multidrug resistance protein 2 (MRP2) or act upon by γ-glutamyl transpeptidase (GGT) to generate reactive platinum–cysteine thiols that can re-enter cells and induce cytotoxicity [11, 15]. This net flux of GSH through platinum conjugation, combined with direct inhibition of glutathione reductase by cisplatin, precipitates a state of reductive stress collapse characterized by GSH depletion and an elevated GSSG:GSH ratio [13]. The resulting pro-oxidant milieu drives Fenton-type reactions and lipid peroxidation chain reactions, yielding MDA and other aldehydic TBARS as end products of PUFA oxidation, which is consistent with the highly significant increase in MDA observed in both the liver and kidney in the current study.

The differential magnitude of cisplatin-induced oxidative injury between the liver and kidney observed in our study, with renal changes being substantially more pronounced across all three parameters, is entirely consistent with the established pharmacokinetics of cisplatin. The kidney concentration of cisplatin is 5–10-fold greater than that of plasma [11, 14]. While the liver is also exposed to high concentrations of cisplatin via portal blood, it exhibits greater phase I and phase II metabolic capacity and higher constitutive antioxidant enzyme expression, rendering it more resistant to acute cisplatin-induced oxidative insult at the dose employed.

The hepatoprotective effects of naringenin observed in the current study attenuate the increase in MDA levels and the preservation of GST activity, with partial maintenance of GSH, which aligns closely with, and substantially extends, the extensive body of literature characterizing naringenin as a potent hepatoprotectant. Hernández-Aquino and Muriel (2018) conducted a comprehensive review of the hepatoprotective mechanisms of naringenin in liver disease models, identifying Nrf2 pathway activation as the primary mechanistic driver: naringenin dissociates Nrf2 from its cytoplasmic repressor Keap1, enabling nuclear translocation and transcriptional upregulation of antioxidant response element (ARE)-driven genes, including those encoding GST, GPx, SOD, catalase, and GR, which collectively amplify the hepatocellular antioxidant shield. This Nrf2/ARE mechanism provides a particularly elegant explanation for the observed preservation of hepatic GST activity in naringenin-treated animals, as GST is a canonical Nrf2 target gene whose expression is directly upregulated by naringenin [17].

Additionally, the hepatoprotective activity of naringenin encompasses the suppression of hepatic inflammation via the inhibition of NF-κB-mediated proinflammatory cytokine production (TNF-α, IL-1β, and IL-6), the downregulation of COX-2 and iNOS, and the attenuation of TGF-β1-mediated hepatic stellate cell activation mechanisms that complement its direct antioxidant effects and are particularly relevant in the context of cisplatin-induced hepatic inflammation [17, 18]. At the lipid membrane level, the intercalation of naringenin into the phospholipid bilayer increases membrane rigidity and reduces fluidity, thereby physically impeding the propagation of lipid peroxidation chain reactions and explaining the observed attenuation of MDA in liver tissue [13]. The lipid-lowering and insulin-sensitizing properties of naringenin further contribute to hepatoprotection by reducing hepatic lipid substrate availability for oxidative attack [31].

The nephroprotective effects of naringenin were more pronounced and statistically robust than its hepatoprotective effects were in the current study, which is consistent with the greater severity of cisplatin-induced renal oxidative stress. The near-complete restoration of renal GSH levels in the cisplatin + naringenin group is particularly noteworthy and suggests that naringenin replenishes renal GSH reserves through at least two complementary pathways: (i) direct antioxidant GSH-sparing by neutralizing ROS before they can oxidize GSH to GSSG, thereby reducing the demand on the GSH/GSSG cycle; and (ii) Nrf2-mediated transcriptional induction of γ-glutamylcysteine synthetase (γ-GCS), the rate-limiting enzyme in de novo GSH biosynthesis [21].

The significant attenuation of renal MDA levels in naringenin-cotreated animals reflects the combined contribution of GSH preservation, membrane-stabilizing effects, and direct radical quenching by the phenolic hydroxyl groups of naringenin. These renal antioxidant effects are consistent with those reported by Al-Dosari et al. (2017), who demonstrated naringenin-mediated attenuation of oxidative stress, apoptosis, and neurotrophic deficits in the diabetic rat retina in a high-oxidative-stress microenvironment mechanistically analogous to the cisplatin-injured proximal tubule [21]. Furthermore, the metal-chelating properties of naringenin conferred by the 3′,4′-catechol-like motif and the 5-OH/4-carbonyl chelation sites of the flavanone skeleton may enable direct sequestration of intracellular platinum ions, reducing the pool of reactive aquated cisplatin species available for GSH conjugation and DNA adduct formation [17].

Of translational importance is the observation by Littest and Schweitzer (1988) that naringenin administration does not reduce renal platinum concentrations, implying that the chemoprotective effects of naringenin are mediated entirely through cytoprotective antioxidant mechanisms rather than by pharmacokinetic interference with cisplatin bioavailability [23]. This is a critical finding for clinical translation: an ideal chemoprotectant should protect healthy organs from cisplatin toxicity without reducing platinum delivery to tumor tissue. In this context, the profile of naringenin is highly favorable, as it can be distinguished from that of glutathione precursors such as N-acetylcysteine, which may theoretically compromise the tumor bioavailability of cisplatin and have yielded inconsistent clinical results [32, 33].

The advantages of naringenin over many related natural chemoprotectant compounds include superior oral bioavailability relative to that of curcumin, established safety in human dietary exposure through citrus consumption, and structural features, particularly its flavanone backbone, that confer unique membrane-intercalating properties not shared by quercetin or thymoquinone [8, 12, 31]. Additionally, naringenin’s documented inhibition of certain CYP450 isoforms (CYP1A1 and CYP2E1) involved in the bioactivation of reactive intermediates may provide an additional layer of xenobiotic chemoprotection beyond glutathione augmentation [22]. Importantly, unlike amifostine, the only FDA-approved cytoprotective agent for the nephrotoxicity of cisplatin, naringenin, is a naturally occurring dietary compound with an excellent safety profile, no known cytoprotective selectivity issues in tumor versus normal tissue, and the potential for oral administration, making it intrinsically more suitable for chronic coadministration with multicycle chemotherapy regimens [10]. Furthermore, naringenin does not alter renal platinum concentrations; therefore, coadministration does not compromise the antineoplastic activity of cisplatin as a prerequisite for clinical translation [23].

The present study constitutes an important proof-of-concept investigation but has several limitations. The biochemical endpoints assessed (MDA, GSH, and GST) represent established but indirect markers of oxidative stress and do not provide direct measurements of the platinum–DNA adduct burden, mitochondrial ROS production, or specific apoptotic signaling cascades. In the future, we have not directly assessed 8-hydroxy-2′-deoxyguanosine (8-OHdG) as a biomarker of oxidative DNA damage, histopathological examination of liver and kidney tissue, serum biochemical markers of kidney and liver function and molecular mechanisms of chemoprotective action by evaluating Nrf2/ARE activation, NF-κB suppression, and mitochondrial protective effects.

## Conclusion

This study demonstrated that oral naringenin coadministration (50 mg/kg body weight/day) significantly attenuated cisplatin-induced oxidative hepatotoxicity and nephrotoxicity in male Swiss albino mice, as evidenced by significant suppression of lipid peroxidation (MDA), near-complete restoration of renal and hepatic GSH levels, and preservation of glutathione S-transferase activity in the liver. The nephroprotective effects were more pronounced than the hepatoprotective effects, which is consistent with the established preferential renal accumulation and toxicity of cisplatin. These findings support a multimechanism model of naringenin-mediated organ protection encompassing direct free-radical scavenging, membrane phospholipid stabilization, Nrf2/ARE-driven antioxidant enzyme induction, and potential metal chelation of reactive platinum species. Naringenin alone had no adverse effects on either organ at the tested dose, confirming its physiological safety profile. Collectively, these data provide a compelling preclinical rationale for further systematic evaluation of naringenin as a safe, orally bioavailable, naturally derived chemoprotectant in cisplatin-based chemotherapy regimens. Future studies integrating histopathological assessment, functional organ markers, tumor efficacy evaluation, mechanistic pathway analysis, and pharmacokinetic characterization will be essential to translate these findings toward clinical application and to establish naringenin as a viable adjunct in oncological supportive care.

## Acknowledgments

The authors gratefully acknowledge the Manipal School of Life Sciences and Manipal Academy of Higher Education, Manipal, for providing support.

## Statements and declarations

### Author Contributions

KDM: Conceptualizations, study design, supervision, data interpretation, manuscript correction and finalization. AD: performed the experiments, performed the analysis and wrote the draft manuscript. All the authors have read and approved the final version of the manuscript.

### Data availability

All the data associated with the study are available within the manuscript. Data will be made available upon request.

